# Anterior cingulate cortex and ventral hippocampal inputs to the basolateral amygdala selectively control generalized fear

**DOI:** 10.1101/620237

**Authors:** Samantha Ortiz, Maeson S. Latsko, Julia L. Fouty, Sohini Dutta, Jordan M. Adkins, Aaron M. Jasnow

**Affiliations:** Department of Psychological Sciences, Kent State University, Kent OH 44242; Brain Health Research Institute, Kent State University, Kent OH 44242

## Abstract

Nearly one third of Americans have been afflicted with an anxiety disorder. A common symptom of anxiety disorders is the over generalization of fear across a broad range of contextual cues. We previously found that the anterior cingulate cortex and ventral hippocampus (vHPC) regulate generalized fear. Here, we investigate the functional projections from the ACC and vHPC to the amygdala and their role in governing generalized fear in a preclinical rodent model. A chemogenetic approach (DREADDs) was used to inhibit glutamatergic projections from the ACC or vHPC that terminate within the basolateral amygdala (BLA) at recent (1 day) or remote (28 days) time points after contextually fear conditioning male mice. Inactivating ACC or vHPC projections to the BLA significantly reduced generalized fear to a novel, nonthreatening context but had no effect on fear to the training context. Further, our data indicate that the ACC-BLA circuit supports generalization in a time-independent manner. We also identified for the first time a strictly time-dependent role of the vHPC-BLA circuit in supporting remote generalized contextual fear. Dysfunctional signaling to the amygdala from the ACC or the hippocampus could underlie over-generalized fear responses that are associated with anxiety disorders. Our findings demonstrate that the ACC and vHPC regulate fear expressed in novel, nonthreatening environments via projections to the BLA but do so as a result of training intensity or time, respectively.

**Significance Statement:** Anxiety disorders are characterized by a common symptom that promotes overgeneralization of fear in non-threatening environments. Dysregulation of the amygdala, anterior cingulate cortex (ACC), or hippocampus has been hypothesized to contribute to increased fear associated with anxiety disorders. Our findings show that the ACC and HPC projections to the basolateral amygdala regulate generalized fear in non-threatening, environments. However, descending ACC projections control fear generalization independent of time, whereas HPC projections play a strictly time-dependent role in regulating generalized fear. Thus, dysfunctional ACC/HPC signaling to the BLA may be a predominant underlying mechanism of non-specific fear associated with anxiety disorders. Our data have important implications for predictions made by theories about aging memories and interactions between the hippocampus and cortical regions.

## Introduction

Exposure to stressful events can precipitate anxiety disorders, which afflict approximately one third of the adult U.S. population (Kessler et al., 2012). A debilitating symptom of many anxiety disorders is the overgeneralization of fear (Dymond et al., 2015; Morey et al., 2015), manifesting as hyperarousal across a range of contexts that are not associated with any aversive event (Lissek et al., 2005; 2010). Moreover, people with anxiety disorders have hyperreactive amygdalae (Shin et al., 2004; 2006) along with decreased anterior cingulate cortex (ACC) (Yamasue et al., 2003; Woodward et al., 2006; Asami et al., 2008; Greenberg et al., 2013) and hippocampal volumes (Gurvits et al., 1996; Shin et al., 2006; Chen and Etkin, 2013). Although these regions are associated with anxiety disorders, there is no evidence demonstrating how these brain areas interact to support over-generalization of fear, leading to the maintenance of anxiety symptomology. In this study, we explore generalized fear – fear occurring in non-threatening contexts – using a preclinical rodent model to identify if glutamatergic projections from the ACC and/or hippocampus to the amygdala regulate generalized fear.

Rodent models of context fear learning have been used for decades to study the underlying mechanisms of fear generalization (for review see Jasnow et al., 2012; 2016; Asok et al., 2018). Twenty-four hours after training mice to fear a context with specific cues, if placed back in the training context, mice display high levels of freezing – a fundamental rodent fear response. If mice are instead placed in a novel context that is different from the training context, they display low levels of freezing, indicating little fear to the novel context. As the time interval between training and testing increases, mice freeze in the novel context at similar levels to those in the training context, generalizing fear to the novel, non-threatening context.

Time-dependent generalized fear is thought to rely on cortical regions (Frankland et al., 2004b; Einarsson et al., 2015), independent of the hippocampus whereas fear responses to specific contexts – *specific fear* – are reliant on the hippocampus (Zola-Morgan and Squire, 1990; Frankland et al., 1998; 2004a; Teyler and Rudy, 2007; Winocur et al., 2007; Wiltgen et al., 2010). We previously identified that generalized fear is simultaneously dependent on the ACC and the ventral hippocampus (vHPC); inactivation of either region reduced fear in a novel, non-threatening context, but left fear to the training context unaltered (Cullen et al., 2015).

Although the ACC and hippocampus are implicated in anxiety disorders (see above citations) and generalized fear (Einarsson and Nader, 2012; Cullen et al., 2015; Zhou et al., 2017), little is known about the circuits through which they govern generalized fear responses. A single study found that circuits connecting the ACC and vHPC in the nucleus reunions are necessary for the learning of specific fear (Xu and Südhof, 2013) – inactivating these circuits prior to training induces rapid fear generalization. However, how the ACC and vHPC outputs govern temporally graded generalized fear during recall is completely unknown. The ACC and vHPC each communicate with the basolateral amygdala (BLA) (Maren and Fanselow, 1995; Cenquizca and Swanson, 2007; Morozov et al., 2011) – a critical region for fear acquisition and expression (Kim and Fanselow, 1992; Kim et al., 1993; Campeau and Davis, 1995; Maren et al., 1996; Schafe et al., 2005; Do-Monte et al., 2016). Thus, we hypothesize that ACC and vHPC projections converge within the BLA to regulate time-dependent contextual generalization of fear.

To identify if ACC and vHPC projections to the BLA regulate generalized fear, we used DREADDs (Armbruster et al., 2007), to selectively express the modified human muscarinic acetylcholine receptor 4 (hM4D) within the ACC or vHPC. We found new evidence that inactivation of ACC or vHPC projections in the BLA dramatically attenuated generalized fear in time-independent and time-dependent processes, respectively; specific fear was unaltered. Our findings suggest that over-generalization of fear in people with anxiety disorders may result from hyperreactive amygdalae due to dysfunctional signaling from the ACC or hippocampus.

## Materials and Methods

### Subjects

Experiments 1 (Fig. 1B), 2 (Fig. 1C-G), 3 (Fig. 2), 5 (Fig. 4), and 6 (Fig. 5) used 224 C57BL/6J male mice. Experiments 4 (Fig 3) and 7 (Fig 6) used 87 F1 male hybrids generated from crossing C57BL/6 males and 129S1SvImJ females (Jackson). All mice were generated from a breeding colony in the Department of Psychological Sciences at Kent State University. Mice were five to seven weeks of age before they were used for experimentation and were group housed (2-5 mice per cage) with free access to food and water in a room maintained on a 12:12 light/dark cycle. All procedures were conducted in a facility accredited by the AALAC, in accordance with the NIH guidelines, and with approval by Kent State University IACUC guidelines.

**Figure 1.**
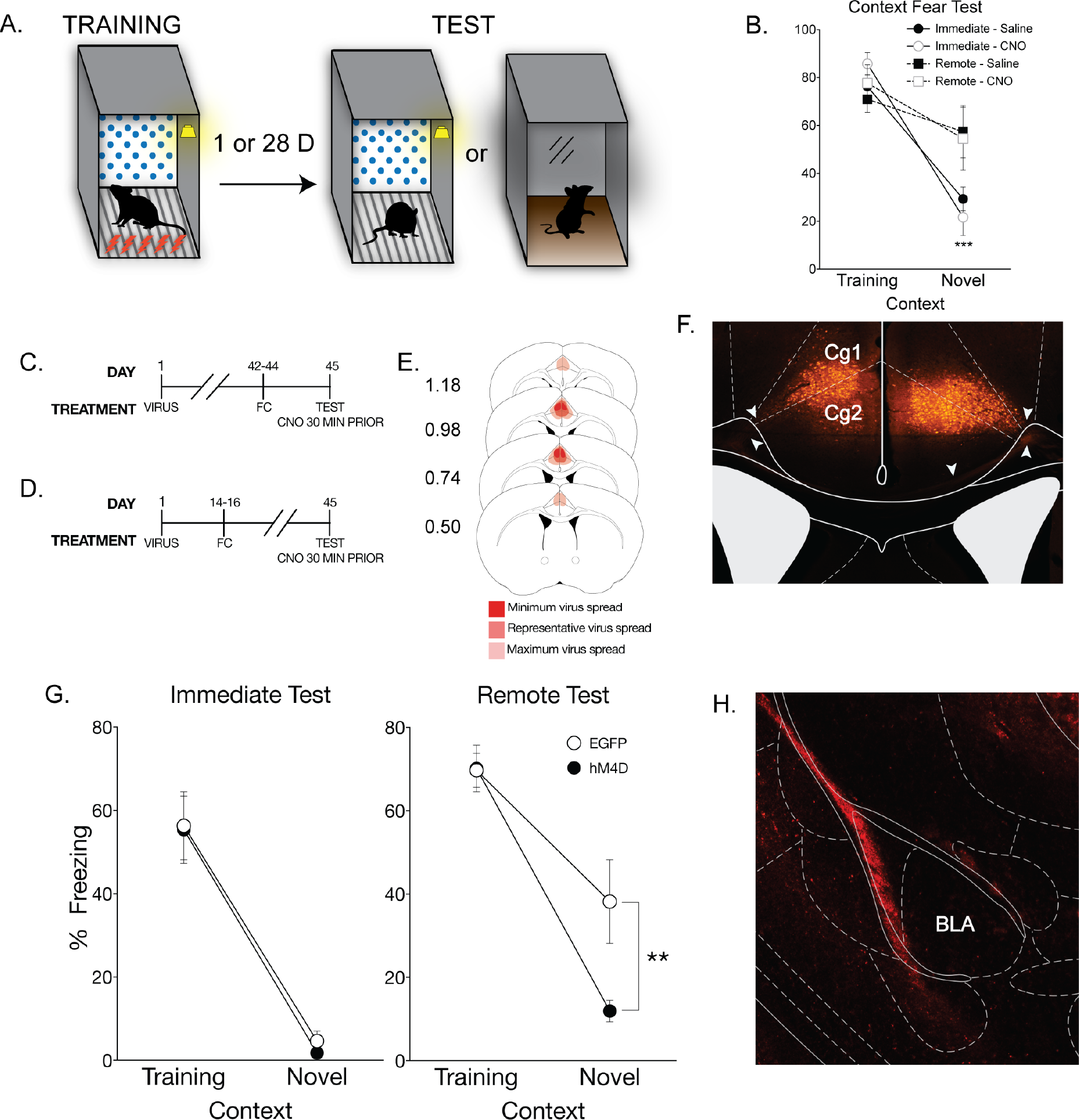
Inactivation of the anterior cingulate cortex eliminates time-dependent generalized context fear. **(A)** All mice underwent context fear conditioning which consisted five unsignaled footshocks (1s, 1.0 mA), each separated by 90s, in the training context which included the conditioning chamber with a polka-dot insert attached to the rear Plexiglas wall, white noise (70db), dim illumination, and the stainless-steel grid floors were cleaned with 70% ethanol. One day or 28 days after training mice were either placed back in the training context or a distinct novel context which included the conditioning chamber with a small exhaust fan, and flat brown Plexiglas floors which were cleaned with 50% Quatricide. There was no visible illumination (illuminated only with an infrared light), and no polka-dot wall insert. **(B)** There was no effect of CNO alone on context dependent fear behavior. As a CNO control experiment, naïve mice were context fear conditioned and given an IP injection of CNO or saline 30 minutes prior to testing either 1 or 28 days after training. Percent freezing levels of animals that received saline (filled symbols) or CNO (open symbols) during immediate (circles) and remote (squares) tests in the training or neutral context were analyzed (± SEM). Two-way ANOVA identified a significant main effect of context at the immediate time point, F_(1,12)_ = 96.40, p< 0.001, but not at the remote time point; mice froze significantly more in the training context than the novel context at 1 but not 28 days after training. ***, significantly different from animals tested in training context, p < 0.001. **(C)** On the first day of the experimental procedures, pAAV-CaMKIIa-hM4D(Gi)-mCherry virus (hM4D) or pAAV-CamKIIa-EGFP (EGFP) was bilaterally infused into the anterior cingulate cortex (ACC). All behavioral tests were completed seven weeks after viral infusions. For the immediate test, mice were tested 1 day after training, **(D)** whereas mice tested at the remote time were tested 28 days after training. All mice were given an IP injection of CNO 30 minutes prior to testing. **(E)** Analysis of transgene expression in all hM4D infusions into the ACC for mice tested with systemic injection of CNO. No expression was observed outside of the ACC for systemic inactivation. Dark red: minimum spread observed and included in analysis; red: representative spread observed; light red: maximum spread observed and included in behavioral analysis. **(F)** Representative image of pAAV-CaMKIIa-hM4D(Gi)-mCherry expression in the ACC. Expression of mCherry was observed throughout the ACC and was typical of a membrane bound fluorophore. White arrows indicate fiber tracts exiting the ACC towards the corpus callosum. (G) hM4D mice administered CNO froze significantly less than EGFP control mice in the novel context only during the remote test, suggesting that inactivation of the ACC eliminates generalized fear at a remote time point. Percent freezing levels of EGFP (◯) and hM4D (⚫) mice during immediate (left panel) and remote (right panel) tests in the training or neutral context were analyzed (± SEM). Two-way ANOVA identified a significant main effect of context at the immediate time point, F_(1,16)_ = 64.2, p< 0.001, and a the remote time point F_(1,17)_ = 52.3, p<.001; mice froze more in the training context than the novel context. However, there was a significant context × treatment interaction only at the remote time point, F_(1,17)_ = 4.64, p< 0.05. *p < 0.05, **p < 0.01, *** < 0.001. **(H)** Representative image of pAAV-CaMKIIa-hM4D(Gi)-mCherry expression in the BLA in a mouse that had virus infused into the ACC. Robust expression of mCherry was observed in the external capsule fibers entering the basolateral amygdala.

**Figure 2.**
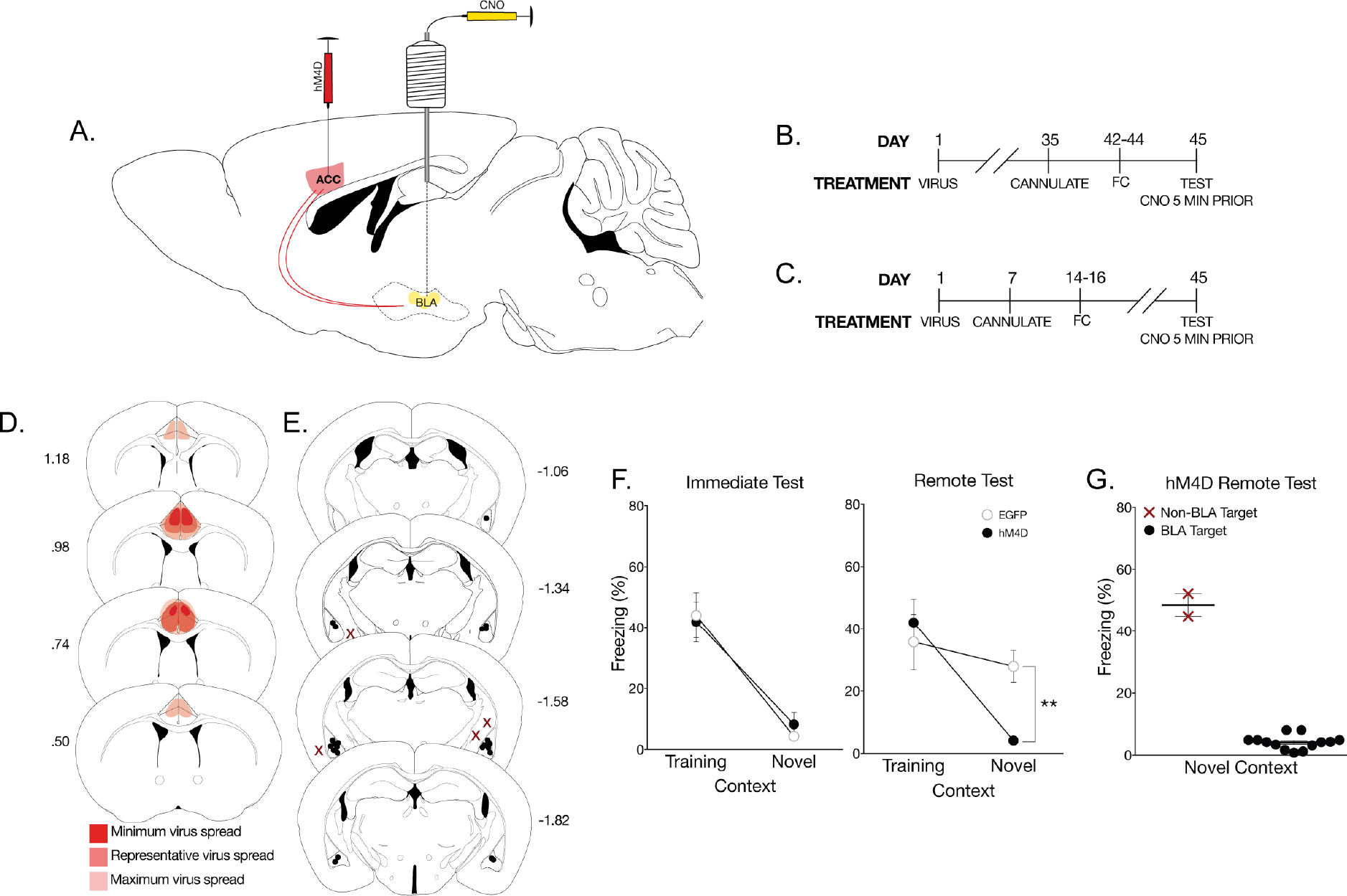
Inactivation of anterior cingulate cortex CamKIIα projections in the basolateral amygdala eliminates time-dependent generalized fear. **(A)** To identify if the ACC regulates fear generalization via CamKIIα projections to the BLA, pAAV-CaMKIIa-hM4D(Gi)-mCherry virus (hM4D) or pAAV-CamKIIa-EGFP (EGFP) was bilaterally infused into the anterior cingulate cortex (ACC) followed by cannulations targeting their axon terminals in the BLA. **(B)** All behavioral tests were completed seven weeks after viral infusions. Cannulations for the BLA were completed one week prior to behavioral training procedures. Mice were tested 1 day or **(C)** 28 days after training. All mice were given a local infusion of CNO into the BLA 5 minutes prior to testing to inactivate ACC CamKIIα projections. **(D)** Analysis of transgene expression in all hM4D mice tested with inactivation of BLA terminals. One mouse was excluded from analysis due to significant hM4D expression in the motor cortex. Dark red: minimum spread observed and included in analysis; red: representative spread observed; light red: maximum spread observed and included in behavioral analysis. **(E)** Cannulation targets within the BLA; black dots indicate animals included in behavioral analyses, red Xs indicate missed targets and used in a site specific control analysis. **(F)** hM4D mice with inactivated CamKIIα projections from the ACC to the BLA froze significantly less than EGFP mice in the novel context, but not in the training context only at the remote test. Percent freezing levels of EGFP (◯) and hM4D (⚫) mice during immediate (left panel) and remote (right panel) tests in the training or neutral context 5 minutes after a microinfusion of CNO were analyzed (± SEM). A two-way ANOVA identified a significant effect of context at the immediate test F_(1,27)_ = 47.1, p< 0.001, and remote test, F_(1,35)_ = 15.6, p< 0.001. As observed previously, there was a significant interaction only at the remote test F_(1,35)_ = 6.71, p< 0.05. Thus, inactivation of ACC CamKIIα projections to the BLA eliminated time-dependent generalized fear. **(G)** hM4D mice with extra-BLA infusions did not show a reduction in freezing in the novel context. Percent freezing levels of hM4D mice tested in the neutral context with missed BLA targeting compared to hM4D mice with specific targeting in the BLA was analyzed (± SEM). A non-parametric Mann-Whitney t-test showed a significant effect of CNO infusion target, p< 0.05. *p < 0.05, **p < 0.01, ***p < 0.001.

**Figure 3.**
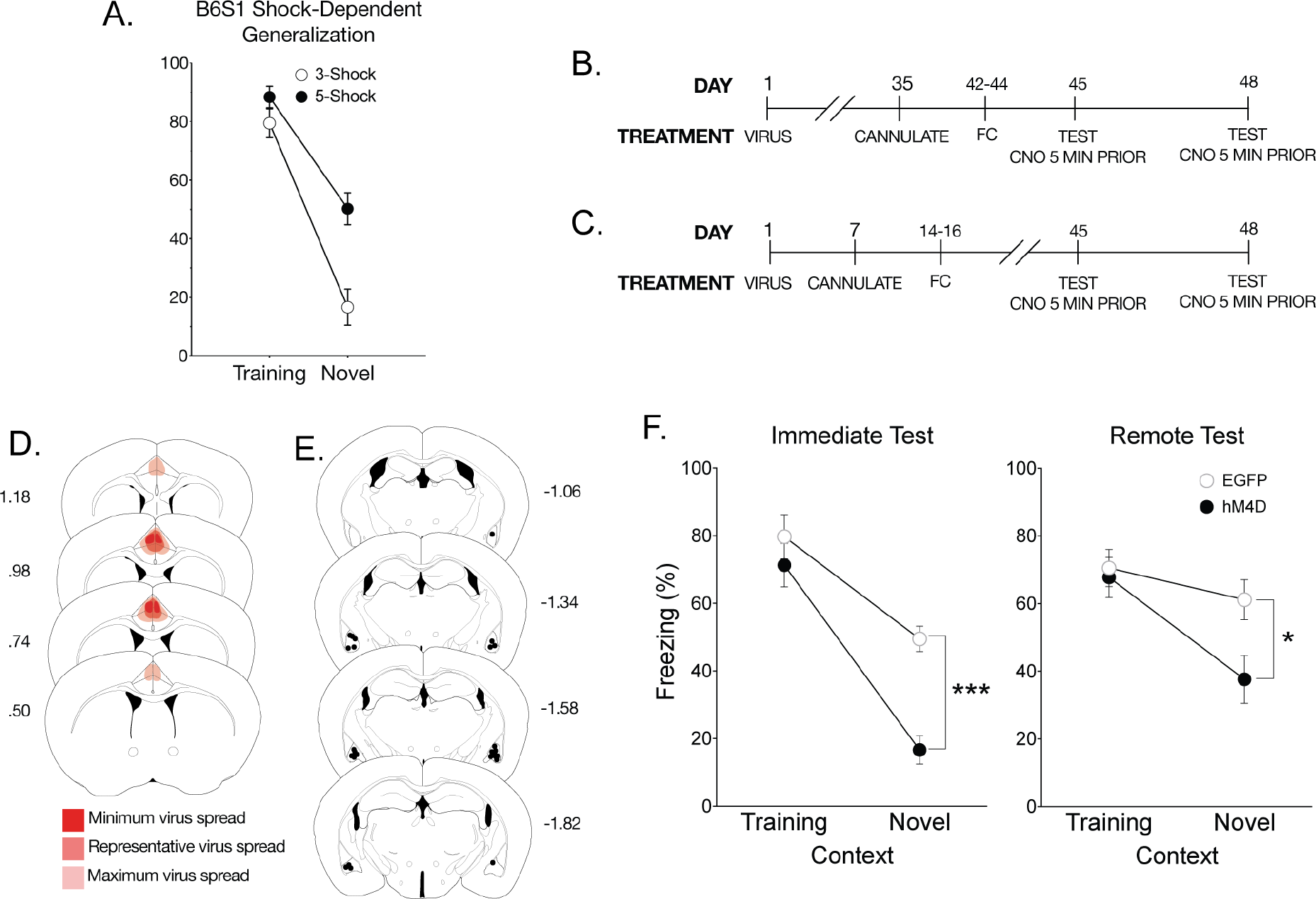
Inactivation of anterior cingulate cortex to basolateral amygdala CamKIIα projections eliminates time-independent generalized fear. **(A)** Hybrid B6S1 mice were tested for contextual fear after training with either 3, 1mA shocks or 5, 1mA shocks. Percent freezing levels of 3 shock (◯) and 5 shock (⚫) trained mice in the training context were analyzed (± SEM). A two-way ANOVA identified significant shock × context interaction F_(1,19)_ = 5.42, p< 0.05, showing that 5-shock training, but not 3-shock training, significantly increased freezing in the novel context at the 24h test. **(B)** All behavioral tests were completed seven weeks after viral infusions. Cannulations for the BLA were completed one week prior to behavioral training procedures. In this experiment immediate generalization was induced using a hybrid mouse line. Mice were tested once in each context at 1 day or **(C)** 28 days after training with a 72-hour inter-test-interval. All mice were given a local infusion of CNO into the BLA 5 minutes prior to testing to inactivate ACC CamKIIα projections. **(D)** As done previously, mice were infused with the hM4D or EGFP virus into the ACC with cannulations targeting the BLA. Viral spread analysis of all hM4D mice tested using a within subject design with inactivation of BLA terminals identified no expression outside of the ACC. Dark red: minimum spread observed and included in analysis; red: representative spread observed; light red: maximum spread observed and included in behavioral analysis. (E) Cannulation targets were analyzed to correct placement into the BLA. No mice had targets localized outside of the BLA in this experiment. **(F)** At recent and remote tests, inactivating CamKIIα projections from the ACC to the BLA significantly reduced freezing to the novel context. Percent freezing levels of EGFP (◯) and hM4D (⚫) mice during within-subject immediate (left panel) or remote (right panel) tests in the training and neutral context 5 minutes after a microinfusion of CNO were analyzed (± SEM). A two-way ANOVA identified significant main effects of context at the immediate F_(1,10)_ = 64.8, p< 0.001 and remote tests F_(1,13)_ = 17.9, p< 0.001. However, for the first time, there was a significant interaction at the immediate F_(1,10)_ = 5.35, p< 0.05 and remote times F_(1,13)_ = 4.93, p< 0.05, suggesting that ACC CamKIIα projections to the BLA control a time-independent form of generalization.

**Figure 4.**
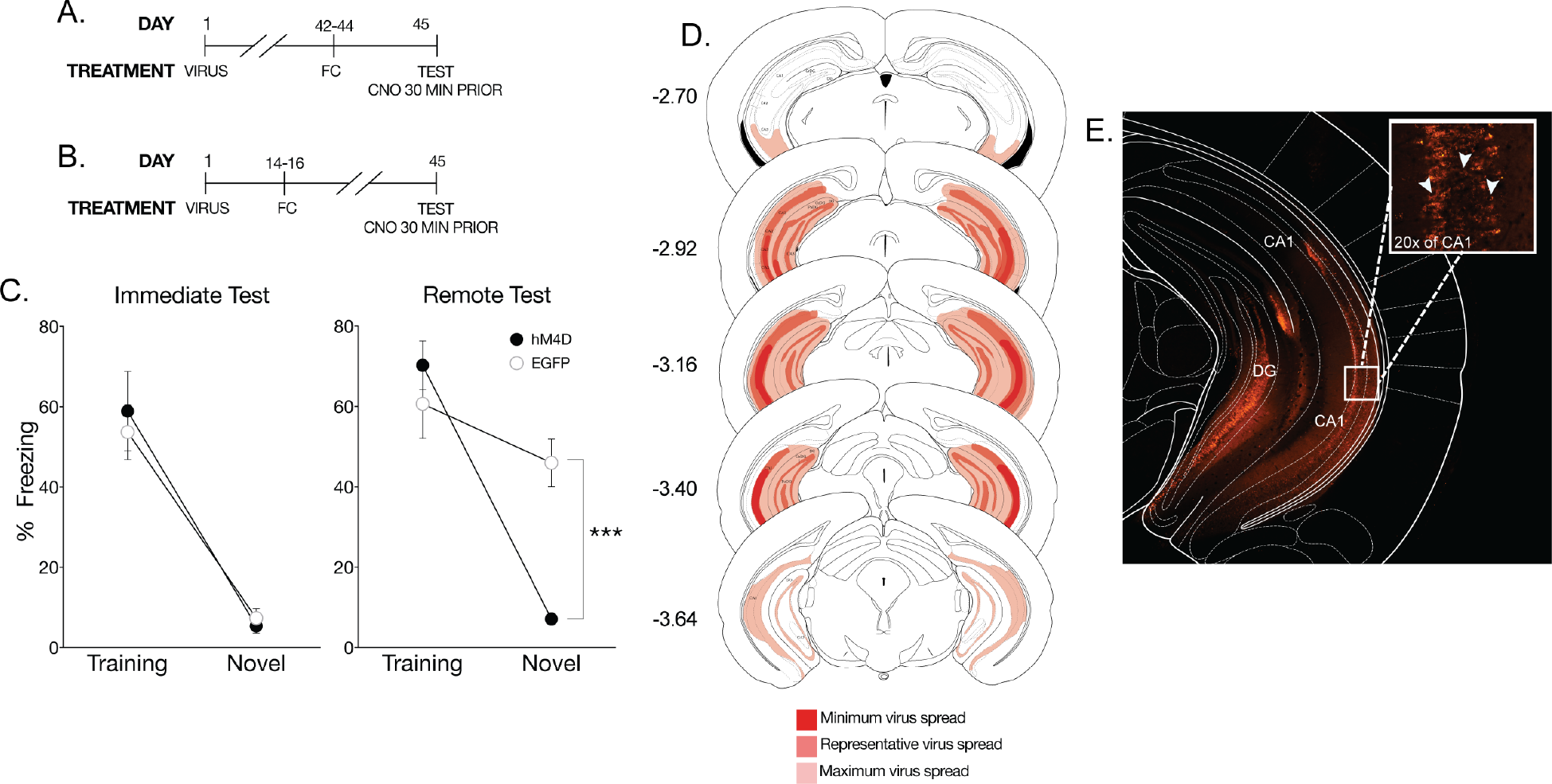
Inactivation of the ventral hippocampus eliminates time-dependent context fear generalization. **(A)** On the first day of the experimental procedures, pAAV-CaMKIIa-hM4D(Gi)-mCherry virus (hM4D) or pAAV-CamKIIa-EGFP (EGFP) was bilaterally infused into the ventral hippocampus (vHPC). All behavioral tests were completed seven weeks after viral infusions. For the immediate test, mice were tested 1 day after training, **(B)** whereas mice tested at the remote time were tested 28 days after training. All mice were given an IP injection of CNO 30 minutes prior to testing. **(C)** Analysis of transgene expression in hM4D infusions into the vHPC for mice tested with systemic injection of CNO. No expression was observed outside of the vHPC. Dark red: minimum spread observed and included in analysis; red: representative spread observed; light red: maximum spread observed and included in behavioral analysis. (D) hM4D mice administered CNO froze significantly less than EGFP control mice in the novel context only. Percent freezing levels of EGFP (◯) and hM4D (⚫) mice during immediate (left panel) and remote (right panel) tests in the training or neutral context were analyzed (± SEM). Two-way ANOVA identified a significant main effect of context at the immediate time point, F_(1,21)_ = 70, p< 0.001, and a the remote time point F_(1,16)_ = 40.9, p<.001; mice froze more in the training context than the novel context. However, there was a significant context × treatment interaction only at the remote time point F_(1,16)_ = 15.9, p< 0.01. *** p < 0.001, suggesting that the vHPC also regulates time-dependent generalized fear. **(E)** Representative photomicrograph of pAAV-CaMKIIa-hM4D(Gi)-mCherry expression in the vHPC. Robust transgene expression was observed throughout the vHPC and typical of a membrane-bound fluorophore. Inset is 20× magnification. White arrows indicate examples of somatic transgene expression.

**Figure 5.**
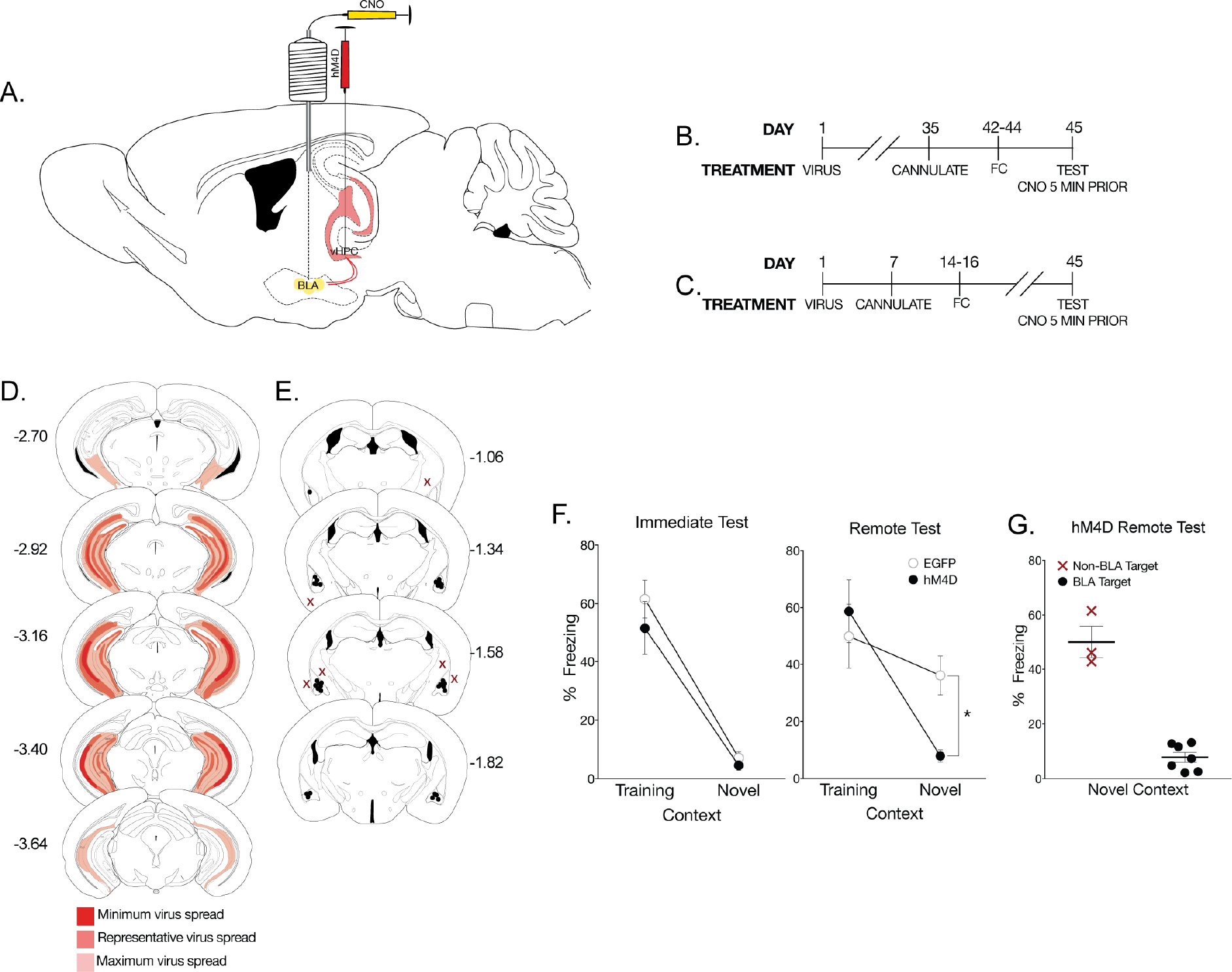
CamKIIα projections from the ventral hippocampus to the basolateral amygdala regulate time-dependent generalized. **(A)** To identify if the vHPC regulates fear generalization via its CamKIIα projections to the BLA, pAAV-CaMKIIa-hM4D(Gi)-mCherry virus (hM4D) or pAAV-CamKIIa-EGFP (EGFP) was bilaterally infused at into the vHPC followed by cannulations targeting the BLA. **(B)** All behavioral tests were completed seven weeks after viral infusions. Cannulations for the BLA were completed one week prior to behavioral training procedures. Mice were tested 1 day or **(C)** 28 days after training. All mice were given a local infusion of CNO into the BLA 5 minutes prior to testing. **(D)**Viral spread analysis of all hM4D mice tested with inactivation of BLA terminals. Dark red: minimum spread observed and included in analysis; red: representative spread observed; light red: maximum spread observed and included in behavioral analysis. **(E)**Cannulation targets within the BLA; black dots indicate animals included in behavioral analyses, red Xs indicate missed targets and used in a site specific control analysis. **(F)**hM4D mice with inactivated CamKIIα projections from the vHPC to the BLA froze significantly less than EGFP mice in the novel context, but not in the training context. Percent freezing levels of EGFP (◯) and hM4D (⚫) mice during immediate (left panel) and remote (right panel) tests in the training or neutral context 5 minutes after a microinfusion of CNO were analyzed (± SEM). A two-way ANOVA identified a significant effect of context at the immediate test F_(1,20)_ = 68.6, p< 0.001, and remote test, F_(1,24)_ = 13.3 p< 0.01. As observed previously, there was a significant interaction only at the remote test F_(1,24)_ = 4.34, p< 0.05. **(G)** hM4D mice with off-target infusions did not show a reduction in freezing in the novel context. Percent freezing levels of hM4D mice tested in the neutral context with missed BLA targeting compared to hM4D mice with specific targeting in the BLA was analyzed (± SEM). A non-parametric Mann-Whitney t-test showed a significant effect of CNO infusion target, p< 0.05. *p < 0.05, **p < 0.01, ***p < 0.001.

**Figure 6.**
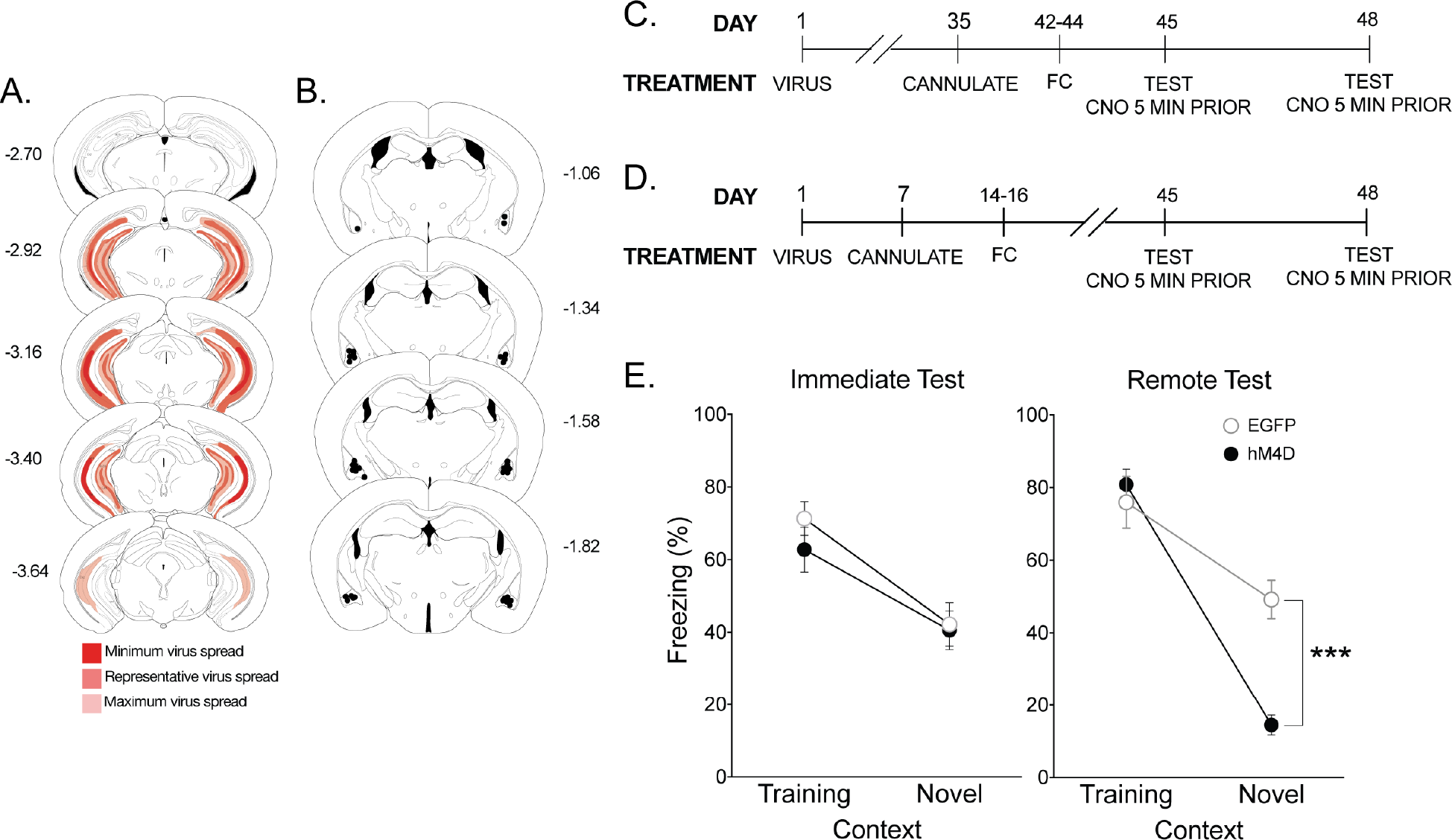
The ventral hippocampus selectively controls time-dependent generalization. **(A)** As done previously, mice were infused with hM4D or EGFP virus into the vHPC with cannulations targeting the BLA. Viral spread analysis of all hM4D mice tested using a within subject design with inactivation of BLA terminals identified no expression outside of the vHPC. Dark red: minimum spread observed and included in analysis; red: representative spread observed; light red: maximum spread observed and included in behavioral analysis. **(B)** Cannulation targets were again analyzed to correct placement into the BLA. There were no missed targets outside of the BLA in this experiment. **(C)** All behavioral tests were completed seven weeks after viral infusions. Cannulations for the BLA were completed one week prior to behavioral training procedures. In this experiment immediate generalization was induced using a hybrid mouse line. Mice were tested once in each context at 1 day or **(D)** 28 days after training with a 72-hour inter-test-interval. All mice were given a microinfusion of CNO into the BLA 5 minutes prior to testing. **(E)** Inactivating CamKIIα projections from the vHPC to the BLA significantly reduced freezing to the novel context only at the remote test. These data suggest that glutamatergic projections from the vHPC to the BLA selectively control time-dependent generalized fear. Percent freezing levels of EGFP (◯) and hM4D (⚫) mice during within-subject immediate (left panel) or remote (right panel) tests in the training and neutral context 5 minutes after a local infusion of CNO were analyzed (± SEM). A two-way ANOVA identified significant main effects of context at the immediate F_(1,13)_ = 19, p< 0.001 and remote tests F_(1,9)_ = 80.9, p< 0.001. After induced generalization, there was a significant interaction only at the remote test F_(1,9)_ = 14.6, p< 0.01. *** p < 0.001.

### Surgical Procedures

Mice were anesthetized with a subcutaneous injection of a Ketamine (75 mg/kg) + Xylazine (10 mg/kg) + Acepromazine (2 mg/kg) cocktail. Following administration of anesthesia, mice were mounted on a stereotaxic apparatus (David Kopf Instruments, Tujunga, CA). The scalp of each mouse was retracted; the skull was adjusted so that bregma and lambda were on the same horizontal plane (within .05mm of each other). Two 0.33 gauge infusion needles were guided to the appropriate coordinates relative to bregma and small bilateral burr holes were drilled. Coordinates for the following brain regions were ACC: .08 mm AP, ±.07 mm ML, −3.6 mm DV from bregma at a 14° angle; vHPC: −3.2 mm AP, ±3.3 mm ML, −4.25 mm DV from bregma. AAV8-CaMKIIa-hM4D(Gi)-mCherry virus (hM4D) (Addgene) or a control virus under the same promoter, AAV8-CamKIIα-EGFP (EGFP) (Addgene) was bilaterally infused at 0.1μL/minute to a total infusion volume of 0.25μL and the needle was left in place for five minutes after completion of the infusion. Upon completion of the virus infusion, the anesthesia was reversed with a subcutaneous injection of atipamezole (0.5 mg/kg).

All behavioral testing was completed seven weeks after viral infusions in order to control for viral expression. Cannulations for the BLA were completed one week prior to behavioral training procedures, controlling for recovery time between the final surgery and the start of behavioral training. Mice were anesthetized and mounted on a stereotaxic apparatus with the same surgical procedures as described above. Two guide cannulae (Plastics One, Roanoke, VA) were surgically implanted bilaterally above the basolateral amygdala (−1.6 mm AP, ±3.4 mm ML, −4.9 mm DV from bregma). Dummy cannulae were inserted into the guide cannulae after surgery. For viral spread analysis and drug targeting for each experiment see figures: 1E,F, and H, 2D-E, 3D-E, 4D, 5D-E, 6A-B.

### Fear Conditioning

Fear conditioning was performed in four identical conditioning chambers (7” W × 7” D × 12”H) containing two Plexiglas walls, two aluminum sidewalls, and a stainless-steel grid-shock floor (Coulbourn Instruments, Allentown, PA). The training context consisted of the conditioning chamber with a polka-dot insert attached to the rear Plexiglas wall, white noise (70db), dim illumination, and the stainless-steel grid floors were cleaned with 70% ethanol. The novel context consisted of the conditioning chamber with no visible illumination (illuminated only with an infrared light), fan, and flat brown Plexiglas floors which were cleaned with 50% Quatricide.

Mice were pre-exposed to the context for five minutes on the two days prior to fear conditioning. Fear conditioning occurred in the training context with five unsignaled footshocks (1s, 1.0 mA), each separated by 90s, mice were removed from the apparatus 30s after the last shock and returned to their home cage. After training, we conducted a 5-minute test in either the training context or the novel context at 24 hours or 28 days after training.

Mice were given 5mg/kg intraperitoneal (IP) injections of clozapine-n-oxide (CNO) (Cayman Chemical) or saline 30 minutes prior to testing in the CNO control experiments and systemic inactivation studies. The dose of 5mg/kg was selected due to common IP injection doses used for DREADD experiments and has shown to have reduced effects on behavior in naïve mice (MacLaren et al., 2016; Jendryka et al., 2019). In experiments in which mice were given a localized infusion of CNO (0.2μL of 650μM at 0.1μL/min), a concentration within the range of those previously reported (Mahler et al., 2014; Vazey and Aston-Jones, 2014; Scofield et al., 2015), the drug was infused five minutes prior to testing in order to inactivate ACC or vHPC projections terminating in the basolateral amygdala. The within-subject fear testing used F1 hybrids in the same training procedures as described previously with counterbalanced testing. F1 hybrids were tested in both the training and novel contexts for five minutes with 72-hours between testing. Five minutes prior to each test, F1 hybrids were given intra-BLA infusions of CNO as previously described.

### Histology

Mice were deeply anaesthetized with pentobarbital sodium and perfused transcardially with 0.9% saline followed by 4% paraformaldehyde. After perfusion, 13.1μL of 0.5% neutral red solution was infused into the guide cannulae for site verification of BLA targets then the brains were extracted. After extraction, brains were post-fixed in 4% paraformaldehyde for 24-hours then transferred to 30% sucrose solution until sectioning. Coronal sections (40μm thick, taken every 120μm) were cut on a freezing microtome, mounted on glass microscope slide, and cover slipped with MOWIOL mounting medium containing 2.5% DABCO before visualization. All imaging was completed on a Nikon Eclipse Ti-S using a Nikon Intensilight C-HGFIE mercury lamp in conjunction with FITC, and Cy3 filters and analyzed using NIS Elements Software. Exclusion criteria for experiments include: unilateral expression of hM4D within the ACC or vHPC or no expression within the vCA1 of the hippocampus. One mouse was excluded due to hM4D cell body expression that significantly exceeded the boundaries of the ACC into the motor cortex. No expression outside of the vHPC was observed.

### Statistical Analyses

Mean freezing during contextual fear testing were analyzed using a 2×2 factorial analysis of variance (ANOVA) on Prism Graphpad statistical software. Statistically significant ANOVAs were followed up with Tukey HSD post hoc comparisons. BLA target comparisons were analyzed using a non-parametric Mann-Whitney t-test on Prism Graphpad. Effect sizes were calculated for completed experiments along with post-hoc power analyses using G*Power 3. Refer to Tables 1–6 for detailed statistical results for each experiment.

**Table 1.** Clozapine-N-Oxide & Hybrid B6S1 Behavior Statistical Summary

| Mouse Strain | Manipulation | Statistical Test | Test Delay | Comparison | F/t Statistic | DF | % Total Variance | P | * | $\eta^2$ | Effect Size | Power | Figure |
| --- | --- | --- | --- | --- | --- | --- | --- | --- | --- | --- | --- | --- | --- |
| C57BL/6 | CNO vs Saline | Two-Way ANOVA | 1 D | Context x Treatment | 2.30 | 1,12 | 2.08 | 0.155 | ns | 0.160 | 0.43 | 0.36 | 1B |
|  |  |  |  | Context | 96.40 | 1,12 | 87.10 | <.001 | *** | 0.889 | 2.85 | 1.00 |  |
|  |  |  |  | Viral Treatment | 0.03 | 1,12 | 0.02 | 0.873 | ns | 0.002 | 0.05 | 0.05 |  |
|  |  |  | 28 D | Context x Treatment | 0.32 | 1,11 | 2.23 | 0.584 | ns | 0.028 | 0.16 | 0.09 |  |
|  |  |  |  | Context | 2.61 | 1,11 | 18.7 | 0.131 | ns | 0.195 | 0.49 | 0.41 |  |
|  |  |  |  | Viral Treatment | 0.80 | 1,11 | 0.606 | 0.774 | ns | 0.007 | 0.08 | 0.06 |  |
|  | Three vs Five Shock |  | 1 D | Context x Treatment | 5.42 | 1,19 | 4.01 | 0.03 | * | 0.222 | 0.53 | 0.68 |  |
|  |  |  |  | Context | 90.60 | 1,19 | 67.10 | <.001 | *** | 0.821 | 2.18 | 1.00 |  |
|  |  |  |  | Shock Treatment | 16.10 | 1,19 | 11.90 | <.001 | *** | 0.78 | 0.29 | 0.26 |  |

## Results

### Clozapine-n-oxide administration in naïve animals has no effect on context fear generalization

Prior to the start of neuronal manipulation with the DREADD system, we tested for non-constitutive effects of CNO on fear generalization. Non-DREADD-infused mice were context fear conditioned and tested in the training context or a distinct novel context where they had not been previously exposed (Fig. 1A) either one or 28 days after training; 30 minutes prior to testing mice were administered CNO or saline. CNO and saline controls displayed high levels of freezing to the training context and significantly lower freezing levels in the novel context at the immediate time point indicating no effect of CNO on normal freezing in either context (Tables 1, 2; Fig. 1B). Furthermore, CNO had no effect on freezing at the remote test; all mice displayed high freezing levels in the training and novel context (Table 1; Fig. 1B). These data indicate that CNO alone, or its potential reverse metabolism to clozapine (Gomez et al., 2017), has no effect on freezing to a specific or generalized context. Thus, any effects observed on fear generalization in the following experiments are due to hM4D receptor inactivation in the targeted region.

**Table 2.** Clozapine-N-Oxide & Hybrid B6S1 Behavior significant post-hoc comparisons summary.

| Statistical Test | Test Delay | Significant Post-Hoc Comparisons<br>Context:Treatment | Mean 1 | Mean 2 | N1 | N2 | t | DF | P | * | Figure |
| --- | --- | --- | --- | --- | --- | --- | --- | --- | --- | --- | --- |
| Pos-Hoc Comparison | 1 D | Training:Saline vs. Novel:Saline | 76.3 | 29.3 | 4 | 4 | 0.8 | 4 | <0.001 | *** | 1B |
|  |  | Training:Saline vs. Novel:CNO | 76.3 | 21.7 | 4 | 4 | 0.8 | 4 | <0.001 | *** |  |
|  |  | Training:CNO vs. Novel:Saline | 85.8 | 29.3 | 4 | 4 | 0.1 | 4 | <0.001 | *** |  |
|  |  | Training:CNO vs. Novel:CNO | 85.8 | 21.7 | 4 | 4 | 0.1 | 4 | <0.001 | *** |  |
| Pos-Hoc Comparison | 1 D | Training:3-Shock vs. Novel:3-Shock | 79.5 | 16.6 | 7 | 6 | 9 | 19 | <.001 | *** |  |
|  |  | Training:3-Shock vs. Novel:5-Shock | 79.5 | 50.2 | 7 | 5 | 4 | 19 | <.001 | *** |  |
|  |  | Training:5-Shock vs. Novel:3-Shock | 88.4 | 16.6 | 5 | 6 | 9.4 | 19 | <.001 | *** |  |
|  |  | Training:5-Shock vs. Novel:5-Shock | 88.4 | 50.2 | 5 | 5 | 4.8 | 19 | <.001 | *** |  |
|  |  | Novel:3-Shock vs. Novel:5-Shock | 16.6 | 50.2 | 6 | 5 | 4.4 | 19 | <.001 | *** |  |

### The anterior cingulate cortex – basolateral amygdala circuit controls time-independent generalized fear

Our initial finding that the ACC plays a critical role in the generalization of context fear (Cullen et al., 2015) was upheld using hM4D inactivation. hM4D-mediated inactivation of the ACC with a systemic injection of CNO eliminated generalized fear to the novel context, but not specific fear to the training context (Tables 3 and 4; Fig. 1G). Therefore, we used the hM4D system with intracranial infusions of CNO to identify the precise ACC circuit that regulates fear generalization. The ACC is known to convey sensory information to the BLA (Morozov et al., 2011; McCullough et al., 2016); therefore, we targeted ACC projection terminals in the BLA.

**Table 3.**
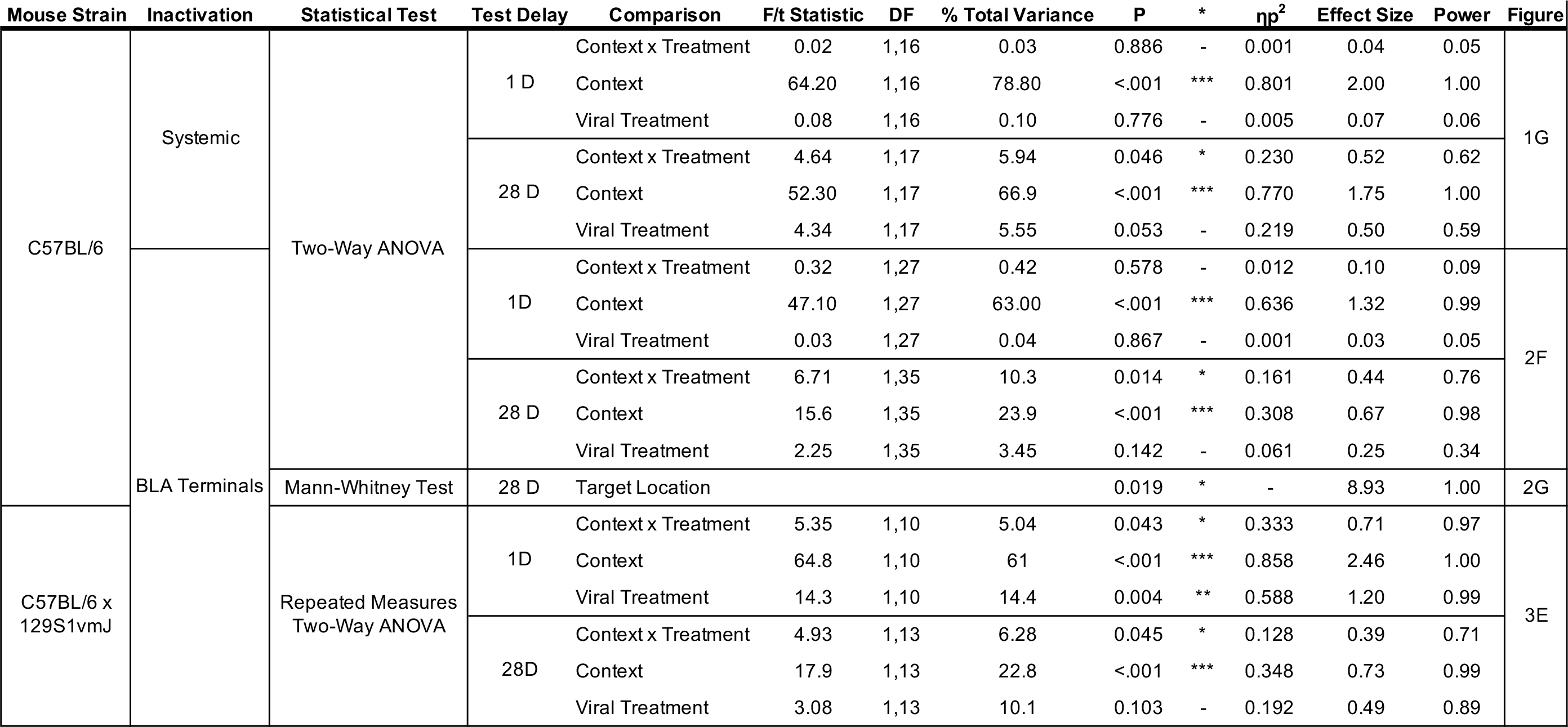
Anterior Cingulate Cortex Statistical Summary

| Mouse Strain | Inactivation | Statistical Test | Test Delay | Comparison | F/t Statistic | DF | % Total Variance | P | * | $\eta p^2$ | Effect Size | Power | Figure |
| --- | --- | --- | --- | --- | --- | --- | --- | --- | --- | --- | --- | --- | --- |
| C57BL/6 | Systemic | Two-Way ANOVA | 1 D | Context x Treatment | 0.02 | 1,16 | 0.03 | 0.886 | - | 0.001 | 0.04 | 0.05 | 1G |
|  |  |  |  | Context | 64.20 | 1,16 | 78.80 | <.001 | *** | 0.801 | 2.00 | 1.00 |  |
|  |  |  |  | Viral Treatment | 0.08 | 1,16 | 0.10 | 0.776 | - | 0.005 | 0.07 | 0.06 |  |
|  |  |  | 28 D | Context x Treatment | 4.64 | 1,17 | 5.94 | 0.046 | * | 0.230 | 0.52 | 0.62 |  |
|  |  |  |  | Context | 52.30 | 1,17 | 66.9 | <.001 | *** | 0.770 | 1.75 | 1.00 |  |
|  |  |  |  | Viral Treatment | 4.34 | 1,17 | 5.55 | 0.053 | - | 0.219 | 0.50 | 0.59 |  |
|  | BLA Terminals | Two-Way ANOVA | 1D | Context x Treatment | 0.32 | 1,27 | 0.42 | 0.578 | - | 0.012 | 0.10 | 0.09 | 2F |
|  |  |  |  | Context | 47.10 | 1,27 | 63.00 | <.001 | *** | 0.636 | 1.32 | 0.99 |  |
|  |  |  |  | Viral Treatment | 0.03 | 1,27 | 0.04 | 0.867 | - | 0.001 | 0.03 | 0.05 |  |
|  |  |  | 28 D | Context x Treatment | 6.71 | 1,35 | 10.3 | 0.014 | * | 0.161 | 0.44 | 0.76 |  |
|  |  |  |  | Context | 15.6 | 1,35 | 23.9 | <.001 | *** | 0.308 | 0.67 | 0.98 |  |
|  |  |  |  | Viral Treatment | 2.25 | 1,35 | 3.45 | 0.142 | - | 0.061 | 0.25 | 0.34 |  |
|  |  | Mann-Whitney Test | 28 D | Target Location |  |  |  | 0.019 | * | - | 8.93 | 1.00 | 2G |
| C57BL/6 x 129S1vmJ |  | Repeated Measures Two-Way ANOVA | 1D | Context x Treatment | 5.35 | 1,10 | 5.04 | 0.043 | * | 0.333 | 0.71 | 0.97 | 3E |
|  |  |  |  | Context | 64.8 | 1,10 | 61 | <.001 | *** | 0.858 | 2.46 | 1.00 |  |
|  |  |  |  | Viral Treatment | 14.3 | 1,10 | 14.4 | 0.004 | ** | 0.588 | 1.20 | 0.99 |  |
|  |  |  | 28D | Context x Treatment | 4.93 | 1,13 | 6.28 | 0.045 | * | 0.128 | 0.39 | 0.71 |  |
|  |  |  |  | Context | 17.9 | 1,13 | 22.8 | <.001 | *** | 0.348 | 0.73 | 0.99 |  |
|  |  |  |  | Viral Treatment | 3.08 | 1,13 | 10.1 | 0.103 | - | 0.192 | 0.49 | 0.89 |  |

**Table 4.** Anterior cingulate cortex significant post-hoc comparisons summary.

| Mouse Strain | Inactivation | Statistical Test | Test Delay | Significant Post-Hoc Comparisons<br>Context:Treatment | Mean 1 | Mean 2 | N1 | N2 | t | DF | P | * | Figure |
| --- | --- | --- | --- | --- | --- | --- | --- | --- | --- | --- | --- | --- | --- |
| C57BL/6 | Systemic | Two-Way ANOVA | 1 D | Training:hM4D vs. Novel:hM4D | 55.4 | 1.79 | 5 | 5 | 5.83 | 16 | <.001 | *** | 1G |
|  |  |  |  | Training:hM4D vs. Novel:EGFP | 55.4 | 4.65 | 5 | 4 | 5.2 | 16 | <.001 | *** |  |
|  |  |  |  | Training:EGFP vs. Novel:hM4D | 56.3 | 1.79 | 6 | 5 | 6.19 | 16 | <.001 | *** |  |
|  |  |  |  | Training:EGFP vs. Novel:EGFP | 56.3 | 4.65 | 6 | 4 | 5.51 | 16 | <.001 | *** |  |
|  |  |  | 28 D | Training:hM4D vs. Novel:hM4D | 70.1 | 11.9 | 6 | 5 | 6.78 | 17 | <.001 | *** |  |
|  |  |  |  | Training:hM4D vs. Novel:EGFP | 70.1 | 38.2 | 6 | 5 | 3.72 | 17 | <.01 | ** |  |
|  |  |  |  | Training:EGFP vs. Novel:hM4D | 69.7 | 11.9 | 5 | 5 | 6.45 | 17 | <.001 | *** |  |
|  |  |  |  | Training:EGFP vs. Novel:EGFP | 69.7 | 38.2 | 5 | 5 | 3.51 | 17 | <.01 | ** |  |
|  |  |  |  | Novel:hM4D vs. Novel:EGFP | 11.9 | 38.2 | 5 | 5 | 2.93 | 17 | <.001 | ** |  |
|  | BLA Terminals | Two-Way ANOVA | 1D | Training:hM4D vs. Novel:hM4D | 41.9 | 8.26 | 7 | 8 | 4.38 | 27 | <.001 | *** | 2F |
|  |  |  |  | Training:hM4D vs. Novel:EGFP | 41.9 | 4.34 | 7 | 8 | 4.89 | 27 | <.001 | *** |  |
|  |  |  |  | Training:EGFP vs. Novel:hM4D | 44 | 8.26 | 8 | 8 | 4.82 | 27 | <.001 | *** |  |
|  |  |  |  | Training:EGFP vs. Novel:EGFP | 44 | 4.34 | 8 | 8 | 5.34 | 27 | <.001 | *** |  |
|  |  |  | 28 D | Training:hM4D vs. Novel:hM4D | 42 | 4.13 | 10 | 13 | 5.08 | 35 | <.001 | *** |  |
|  |  |  |  | Training:EGFP vs. Novel:hM4D | 35.7 | 4.13 | 8 | 13 | 3.96 | 35 | <.001 | *** |  |
| C57BL/6 x<br>129S1vmJ | BLA Terminals | Repeated Measures<br>Two-Way ANOVA | 1D | hM4D: Training vs. Novel | 71.3 | 16.7 | 6 | 6 | 7.33 | 10 | <.001 | *** | 3E |
|  |  |  |  | EGFP: Training vs. Novel | 79.7 | 49.5 | 6 | 6 | 4.05 | 10 | <.01 | ** |  |
|  |  |  |  | Novel: hM4D vs. EGFP | 16.7 | 49.5 | 6 | 6 | 4.32 | 20 | <.001 | *** |  |
|  |  |  | 28D | hM4D: Training vs. Novel | 67.7 | 37.6 | 8 | 8 | 4.72 | 13 | <.001 | *** |  |
|  |  |  |  | Novel: hM4D vs. EGFP | 37.6 | 61.1 | 8 | 7 | 2.66 | 26 | <.05 | * |  |

Mice with hM4D or EGFP virus in the ACC were context fear conditioned; five minutes prior to testing all mice were administered intracranial infusions of CNO via guide cannulae into the BLA (Fig 2A-C). Inactivation of the hM4D-expressing terminals from the ACC in the BLA did not affect freezing in the training or novel context during the immediate test, both hM4D and EGFP groups displayed high freezing in the training context and low freezing in the novel context (Tables 3 and 4; Fig 2F, left panel). However, inactivating ACC terminals in the BLA significantly reduced freezing only in the novel context 28 days after training (Fig. 2F, right panel), whereas EGFP mice displayed equivalent freezing in the training and novel contexts – indicating generalized fear. The reduction of fear generalization in hM4D mice was specific to terminal inactivation within the BLA; hM4D mice with extra-BLA infusions froze significantly more in the novel context than those with intra-BLA infusions (Table 3; Fig. 2G). Thus, we established that projections from the ACC to the BLA are critical for promoting generalized fear at remote testing points.

Are the ACC projections to the BLA that support generalized fear restricted solely to remote tests? If generalization occurs rapidly, does the ACC-BLA circuit still control generalization? Based on our previous findings (Cullen et al., 2015) and the experiments above, we predicted that ACC projections to the BLA would only support generalized fear that develops over time. In the third experiment we used the F1 hybrids of C57BL/6J crossed with 129S1/SvImJ – a hybrid mouse line used by several laboratories to study mechanisms of contextual fear (Frankland et al., 2004b; Smith et al., 2007; Wiltgen and Silva, 2007; Wiltgen et al., 2010; Tanaka et al., 2014) due to their rapid learning and high reliability in fear learning. This gave us the advantage of ensuring that our experimental results were not restricted to C57BL/6J mice, as there is considerable variability in learning and behavior across mouse lines (Hefner et al., 2008). We first performed behavioral parametrics with the F1 hybrid line and found a significant effect of number of shocks on the timing of generalization (Table 1; Fig. 3A). Hybrid mice displayed high levels of freezing in the novel context one day after training if the mice received five footshocks, yet this was not observed if the mice received only three footshocks (Table 1; Fig. 3A); thus, providing a novel opportunity to study the role of the ACC-BLA-vHPC circuit in non-temporally graded generalization.

Experimental procedures were carried out as described in experiment two; however, mice were tested a second time 72-hours after the first test in the opposite context to reduce potential testing-order effects and allow for CNO to be completely metabolized before the second test (Fig. 3B and C). Hybrid mice with EGFP virus displayed increased freezing in the novel context during immediate and remote tests (Fig 3F, left panel). Unexpectedly, hM4D inactivation of the projections from the ACC to the BLA at both the immediate and remote tests reduced freezing in the novel context but not in the training context (Table 3 and 4; Fig. 3F), indicating that projections from the ACC to the BLA promote freezing to a novel context in a time-independent manner. The ACC-BLA pathway controls generalized fear to the novel context but not specific fear to the training context; this effect is upheld across mouse strains and experimental testing designs.

### The ventral hippocampus - basolateral amygdala circuit controls time-dependent generalized fear

In addition to identifying the ACC as a critical locus supporting generalized contextual fear, we previously identified that the vCA1 of the hippocampus also underlies generalized contextual fear at remote time points (Cullen et al., 2015). This finding was replicated by using hM4D to inactivate the vHPC. Inactivation of the vHPC with a systemic injection of CNO significantly reduced generalized fear to the novel context but not specific fear to the training context at a remote time point (Tables 5 and 6; Fig. 4C). As done with Experiment 2, we used intracranial infusions of CNO to identify the vHPC circuit that regulates fear generalization. Given that the vCA1 of the hippocampus has direct connections with the BLA (Cenquizca and Swanson, 2007; Fanselow and Dong, 2010) and is thought to be crucial for conveying contextual information to the BLA (Maren and Fanselow, 1995; Huff et al., 2016) we targeted vHPC projections terminating in this region.

**Table 5.** Ventral hippocampus cortex statistical analysis summary.

| Mouse Strain | Inactivation | Statistical Test | Test Delta | Comparison | F/t Statistic | DF | % Total Variance | P | * $\eta^2$ | Effect Size | Power | Figure |
| --- | --- | --- | --- | --- | --- | --- | --- | --- | --- | --- | --- | --- |
| C57BL/6 | Systemic | Two-Way ANOVA | 1 D | Context x Treatment | 0.36 | 1,21 | 0.40 | 0.553 | - | 0.13 | 0.09 | 4F |
|  |  |  |  | Context | 70.00 | 1,21 | 76.20 | <.001 | *** | 1.82 | 1.00 |  |
|  |  |  |  | Viral Treatment | 0.07 | 1,21 | 0.08 | 0.79 | - | 0.06 | 0.06 |  |
|  |  |  | 28 D | Context x Treatment | 15.90 | 1,16 | 20.2 | 0.001 | ** | 1.00 | 0.97 |  |
|  |  |  |  | Context | 40.90 | 1,16 | 52.1 | <.001 | *** | 1.60 | 0.99 |  |
|  |  |  |  | Viral Treatment | 5.79 | 1,16 | 7.38 | 0.029 | * | 0.60 | 0.71 |  |
|  | BLA Terminals | Two-Way ANOVA | 1D | Context x Treatment | 0.36 | 1,20 | 0.39 | 0.556 | - | 0.13 | 0.10 | 5F |
|  |  |  |  | Context | 68.60 | 1,20 | 75.10 | <.001 | *** | 1.85 | 1.00 |  |
|  |  |  |  | Viral Treatment | 1.05 | 1,20 | 1.15 | 0.318 | - | 0.22 | 0.19 |  |
|  |  |  | 28 D | Context x Treatment | 4.34 | 1,24 | 10.5 | 0.048 | * | 0.43 | 0.51 |  |
|  |  |  |  | Context | 13.3 | 1,24 | 32.2 | 0.001 | ** | 0.76 | 0.94 |  |
|  |  |  |  | Viral Treatment | 1.21 | 1,24 | 2.92 | 0.283 | ns | 0.22 | 0.18 |  |
|  |  | Mann-Whitney Test | 28 D | Target Location |  |  |  | 0.017 | * | 4.02 | 0.99 | 5G |
| C57BL/6 x 129S1vmJ |  | Repeated Measures Two-Way ANOVA | 1D | Context x Treatment | 0.348 | 1,13 | 0.798 | 0.565 | ns | 0.16 | 0.19 | 6E |
|  |  |  |  | Context | 19 | 1,13 | 43.5 | <.001 | *** | 1.21 | 1.00 |  |
|  |  |  |  | Viral Treatment | 0.952 | 1,13 | 1.71 | 0.347 | ns | 0.24 | 0.35 |  |
|  |  |  | 28D | Context x Treatment | 14.6 | 1,9 | 12.4 | 0.004 | ** | 1.17 | 0.99 |  |
|  |  |  |  | Context | 80.9 | 1,9 | 68.6 | <.001 | *** | 2.75 | 1.00 |  |
|  |  |  |  | Viral Treatment | 6.95 | 1,9 | 7.02 | 0.027 | * | 0.88 | 0.99 |  |

**Table 6.** Ventral hippocampus significant post-hoc comparisons summary.

| Mouse Strain | Inactivation | Statistical Test | Test Delay | Significant Post-Hoc Comparisons<br>Context:Treatment | Mean 1 | Mean 2 | N1 | N2 | t | DF | P | * | Figure |
| --- | --- | --- | --- | --- | --- | --- | --- | --- | --- | --- | --- | --- | --- |
| C57BL/6 | Systemic | Two-Way ANOVA | 1 D | Training:hM4D vs. Novel:hM4D | 58.9 | 5.36 | 6 | 7 | 6.5 | 21 | <.001 | *** | 4F |
|  |  |  |  | Training:hM4D vs. Novel:EGFP | 58.9 | 7.35 | 6 | 6 | 6 | 21 | <.001 | *** |  |
|  |  |  |  | Training:EGFP vs. Novel:hM4D | 53.7 | 5.36 | 6 | 7 | 5.8 | 21 | <.001 | *** |  |
|  |  |  |  | Training:EGFP vs. Novel:EGFP | 53.7 | 7.35 | 6 | 6 | 5.4 | 21 | <.001 | *** |  |
|  |  |  | 28 D | Training:hM4D vs. Novel:hM4D | 70.2 | 7.1 | 5 | 5 | 7.3 | 16 | <.001 | *** |  |
|  |  |  |  | Training:hM4D vs. Novel:EGFP | 70.2 | 46 | 5 | 5 | 2.8 | 16 | < 0.05 | * |  |
|  |  |  |  | Training:EGFP vs. Novel:hM4D | 60.7 | 7.1 | 5 | 5 | 6.2 | 16 | <.001 | *** |  |
|  |  |  |  | Novel:hM4D vs. Novel:EGFP | 7.1 | 46 | 5 | 5 | 4.5 | 16 | <.001 | *** |  |
|  | BLA Terminals | Two-Way ANOVA | 1D | Training:hM4D vs. Novel:hM4D | 51.6 | 4.57 | 6 | 5 | 5.2 | 20 | <.001 | *** | 5F |
|  |  |  |  | Training:hM4D vs. Novel:EGFP | 51.6 | 7.16 | 6 | 6 | 5.2 | 20 | <.001 | *** |  |
|  |  |  |  | Training:EGFP vs. Novel:hM4D | 61.6 | 4.57 | 7 | 5 | 6.5 | 20 | <.001 | *** |  |
|  |  |  |  | Training:EGFP vs. Novel:EGFP | 61.6 | 7.16 | 7 | 6 | 6.6 | 20 | <.001 | *** |  |
|  |  |  | 28 D | Training:hM4D vs. Novel:hM4D | 58.7 | 7.9 | 7 | 6 | 3.9 | 24 | < .001 | *** |  |
|  |  |  |  | Training:EGFP vs. Novel:hM4D | 50 | 7.9 | 7 | 6 | 3.2 | 24 | <.01 | ** |  |
|  |  |  |  | Novel:hM4D vs. Novel:EGFP | 7.9 | 36.1 | 6 | 8 | 2.2 | 24 | <.05 | * |  |
| C57BL/6 x 129S1vmJ |  | Repeated Measures<br>Two-Way ANOVA | 1D | hM4D: Training vs. Novel | 62.8 | 40.6 | 7 | 7 | 2.6 | 13 | <.05 | * | 6E |
|  |  |  |  | EGFP: Training vs. Novel | 71.3 | 42.2 | 8 | 8 | 3.6 | 13 | <.01 | ** |  |
|  |  |  | 28D | hM4D: Training vs. Novel | 80.8 | 14.5 | 5 | 5 | 8.7 | 9 | <.001 | *** |  |
|  |  |  |  | EGFP: Training vs. Novel | 76 | 49.2 | 6 | 6 | 3.8 | 9 | <.01 | ** |  |
|  |  |  |  | Novel:hM4D vs. Novel:EGFP | 14.5 | 49.2 | 5 | 6 | 4.5 | 18 | <.001 | *** |  |

Mice with hM4D virus or EGFP control virus in the vHPC were context fear conditioned; five minutes prior to testing all mice were given intracranial infusions of CNO via guide cannulae into the BLA (Fig. 5A-C). Inactivation of hM4D terminals from the vHPC in the BLA did not affect freezing in the training or novel context during the immediate test; both hM4D and EGFP groups displayed high freezing in the training context and low freezing in the novel context (Tables 5 and 6; Fig. 5F, left panel). When mice were tested 28 days after training, EGFP-expressing mice displayed equivalent freezing levels in the training and novel contexts (Tables 5 and 6; Fig. 5F, right panel) – indicating generalized fear. hM4D inactivation of the vHPC terminals in the BLA significantly reduced freezing in the novel context but did not alter freezing in the training context. Again, this effect observed in hM4D-expressing mice was specific to projections from the vHPC terminating in the BLA. HM4D mice with targets outside of the BLA froze significantly more in the novel context at a remote time point than those with correct target placement within the BLA even though they both expressed hM4D and received intracranial CNO infusions (Table 5; Fig. 5G). These findings indicate that activity of vHPC projections – likely via vCA1 outputs (Cenquizca and Swanson, 2007; Cullen et al., 2015) – to the BLA promote generalized fear, but only at a remote time point.

Are the vHPC projections to the BLA that support generalized fear restricted to remote tests? As with Experiment 3, during the immediate test EGFP F1 hybrids displayed increased freezing in the novel context (Tables 5 and 6; Fig. 6E, left panel) – displaying immediate fear generalization. However, unlike the results from ACC-BLA circuit, inactivation of vHPC terminals in the BLA at the immediate time point did not reduce freezing in the novel context nor the training context; reduced generalization was only observed at the remote time point (Tables 5 and 6; Fig. 6E). Given that our previous tests in the novel context at the immediate time point had a floor effect, these experiments identified for the first time a strictly time-dependent role of the vHPC-BLA circuit in supporting generalized contextual fear. Conversely, the ACC governs generalization at both recent and remote tests. Thus, our evidence supports a role for the ACC in supporting generalized fear regardless of the passage of time, whereas the vHPC is engaged in support of generalized fear only at a remote time point.

## Discussion

Clinical studies implicate that the hyperreactive amygdalae observed in people with anxiety disorders may be due to an inhibitory dysregulation caused by a malfunctioning anterior cingulate cortex and hippocampus (Gurvits et al., 1996; Yamasue et al., 2003; Shin et al., 2006; Woodward et al., 2006; Asami et al., 2008; Chen and Etkin, 2013; Greenberg et al., 2013). These studies are limited in making causal conclusions about connectivity, as they associate hyperactive amygdalae with decreased volume and activity of the ACC or hippocampus. Here, we identified causal relationships that fear to novel contexts is in fact regulated by the glutamatergic, CamKIIα-expressing projection neurons from the ACC and vHPC to the BLA but via separate training and time-dependent mechanisms. The regulation of generalized fear by projections from the ACC to the BLA is a time-independent effect that may depend on the strength of the training based on our finding that 5-shock, not 3-shock, training induced immediate generalization. These findings support recent hypotheses that propose that the ACC regulates generalized fear responses (Teyler and Rudy, 2007; Winocur et al., 2007; Einarsson and Nader, 2012; Cullen et al., 2015), but not specific fear responses. The time-independent mechanism of the ACC-BLA connection is in contrast to what we observed with the vHPC. When we induced rapid generalization, inactivation of projections from the vHPC to the BLA did not reduce freezing in the novel context. Generalization was only eliminated when the vHPC-BLA circuit was inactivated at a remote time point. Thus, the vHPC-BLA circuit plays a specific role in time-*dependent* generalization of contextual fear.

We have consistently observed a role for the ACC that is specific to generalized fear responding (Cullen et al., 2015), and this is supported by other recent work (Einarsson et al., 2015). We note two prior studies which found that the ACC regulates specific fear responses at remote time points after training (Frankland et al., 2004a; Goshen et al., 2011). In one case, this discrepancy could be due to specific methodological differences during testing; we performed local intracranial infusions of CNO without anesthetizing mice prior to testing unlike the previous study (Frankland et al., 2004a). In the other case, the authors performed tone-dependent fear training with context as background and used multiple recall tests in the same context (Goshen et al., 2011). Here, we used unsignaled shocks to train specifically for contextual fear and mice were only tested in a single context once. This discrepancy provides evidence that ACC regulation of fear responses is related to the strength – and type – of the fear training. This was not the case for the role of the vHPC in generalized fear responding.

For decades, the focus of identifying neural mechanisms of fear responding has been the dorsal hippocampus (dHPC), and much of the current theory is based on experiments within this region (Squire and Alvarez, 1995a; Frankland et al., 1998; Teyler and Rudy, 2007; Winocur et al., 2007; Wiltgen et al., 2010; Hardt et al., 2013; Winocur et al., 2013). Notably, the experiments described here, and our previous study (Cullen et al., 2015), are the only studies to date examining the role of vHPC in generalized fear responses. Generalized, remote fear responses require the vHPC; whereas the dHPC is crucial for maintaining specific fear responses (Frankland et al., 1998; Wiltgen et al., 2010; Winocur et al., 2013; Cullen et al., 2015). Over time, activity of the vHPC and its projections to the BLA exert greater control over generalized fear rather than maintaining control over specific fear, like the dHPC. Our vHPC results also emphasize that there is a dissociation between the roles of the ventral and dorsal hippocampus in the control of fear processing, an effect that has support from neuroanatomical and connectivity studies (Fanselow and Dong, 2010), but limited systems and behavioral support (Morris, 1981; Maren and Holt, 2004; Hunsaker and Kesner, 2008). The present data also have important implications for predictions that are made by theories about aging fear memories and interactions between the hippocampus and cortical regions (Squire and Alvarez, 1995a; Teyler and Rudy, 2007; Winocur et al., 2007; Hardt et al., 2013).

Systems consolidation hypothesizes that memories stored in the neocortex are identical to those encoded by the hippocampus and does not address time-dependent changes in memory specificity (Squire and Alvarez, 1995a). Our previous (Cullen et al., 2015) and current findings challenge the view that neocortical stored memories are identical to those stored in the hippocampus. In addition, our data suggest that aged memories continue to be dependent on the hippocampus, albeit control shifts to the ventral region. Another memory hypothesis suggests that specific memories are initially dependent on the hippocampus and are transformed to schematic – generalized – memories as they are stored in the neocortex, called the transformation hypothesis (Winocur et al., 2007; 2013), which stems from Multiple Trace Theory (Nadel and Moscovitch, 1997). In the transformation hypothesis, both the schematic memory and the specific memory are continuously accessible; however, specific memories are *always* dependent on the hippocampus whereas generalized memories are dependent on the neocortex as they are transformed over time– independent of the hippocampus. Therefore, at remote time points there can be two memory traces and either can be accessed depending on the situational requirements.

Our data challenge the transformation hypothesis’ notion that neocortical regions control generalized memories as a function of the training-to-testing interval; our data here show that memories may be immediately stored in a generalized state within the ACC. Experiments employing immediate post-training inactivation of the ACC followed by a test for generalization within a novel context are needed in order to confirm the immediate storage hypothesis. Thus, our current data support the neocortex’s involvement in generalized memories, but not that generalized memories are transformed over time, or that they are independent of the hippocampus.

Studies in full support of the transformation hypothesis thus far have not found evidence for a functional dissociation between the dorsal and ventral hippocampus on generalization (Winocur et al., 2007; 2009), suggesting that the hippocampus - as a whole - is not required for generalized memory recall. Here, we discovered that immediately generalized memories do not require the vHPC, whereas remote generalized memories do – showing an opposite role of that of the dHPC. Thus, our data, in combination with recent findings (Lynch et al., 2017; Zhou et al., 2017), suggest that transformation of a specific fear memory into a generalized form may actually involve a shift in control over memory recall from the dHPC to the vHPC over time.

Utilizing chemogenetics, we reliably replicated the effects of the ACC and vHPC regulating fear generalization via projections to the BLA; however, there have been recent validity threats to the DREADD system. The DREADD activator – CNO – may be reverse metabolized into clozapine with widespread effects and non-specific binding of the DREADD receptor (MacLaren et al., 2016; Whissell et al., 2016; Gomez et al., 2017; Manvich et al., 2018). To control for potential off-target effects of CNO, we fear conditioned naïve mice and tested them 30 minutes after an injection of CNO or saline. We found no effect of CNO on contextual fear or the generalization of contextual fear – eliminating the potential confound of CNO specifically for our paradigm. Additionally, intracranial infusions of CNO directly into the BLA replicated the systemic DREADD inactivation findings, and mice expressing hM4D with targets outside the BLA displayed normal freezing behavior in the novel context. Although one study reported off target effects with lower a concentration of CNO when locally infused near the hypothalamus (Stachniak et al., 2014), the small volume of the infusions used here (0.2µL), and the lack of any behavioral effect when CNO was infused outside of the BLA strongly suggests that our observed results were not due to off target effects of CNO – or its reversal into clozapine – and that the effects were specific to inactivation of axonal projections terminating in the BLA. A few reports suggest that CNO must be first converted into clozapine in order to cross the blood brain barrier and exert its effects (Bender et al., 1994; Gomez et al., 2017). Our intra-BLA infusions surpass the blood-brain barrier, therefore CNO – not clozapine – in experiments 3, 4, 6, and 7 specifically acted on the DREADD receptors in virally infused mice.

These findings help to uncover part of the neural connectome involved in both specific and general fear responses, which is critical for understanding how humans and non-humans alike express fearful responses in safe environments (pathological generalization). Clinical research hypothesizes that reduced volume of the ACC and hippocampus restricts normal inhibitory function on the amygdala leading to increased fear responding (Gurvits et al., 1996; Schuff et al., 2001; Yamasue et al., 2003; Shin et al., 2006; Woodward et al., 2006; Asami et al., 2008; Chen and Etkin, 2013; Greenberg et al., 2013). Our findings confirm that the ACC and hippocampus, specifically the vHPC, regulate fear in novel, or non-threatening, environments through their outputs to the amygdala. Furthermore, they do so in a functionally different manner. The ACC time-independently controls generalization, whereas the vHPC plays a strictly time-dependent role in regulating generalized fear. Thus, dysfunctional signaling to the BLA from the ACC or hippocampus – in combination or alone – may be a predominant underlying mechanism of non-specific fear responses associated with anxiety disorders in clinical populations.

## Acknowledgements

We acknowledge the intellectual contributions of Dr. David Riccio and Dr. John Johnson. We also acknowledge the research assistance of Courtney Costanzo and Jasmin Beaver in the data collection. We also acknowledge the contributions of the animal care staff in the Department of Psychological Sciences.

## References

Armbruster BN, Li X, Pausch MH, Herlitze S, Roth BL (2007) Evolving the Lock to Fit the Key to Create a Family of G Protein-Coupled Receptors Potently Activated by an Inert Ligand. Proc Natl Acad Sci USA 104:5163–5168.

Asami T, Hayano F, Nakamura M, Yamasue H, Uehara K, Otsuka T, Roppongi T, Nihashi N, Inoue T, Hirayasu Y (2008) Anterior cingulate cortex volume reduction in patients with panic disorder. Psychiatry and Clinical Neurosciences 62:322–330.

Asok A, Kandel ER, Rayman JB (2018) The Neurobiology of Fear Generalization. Front Behav Neurosci 12:329.

Bender D, Holschbach M, Stöcklin G (1994) Synthesis of n.c.a. carbon-11 labelled clozapine and its major metabolite clozapine-N-oxide and comparison of their biodistribution in mice. Nucl Med Biol 21:921–925.

Campeau S, Davis M (1995) Involvement of subcortical and cortical afferents to the lateral nucleus of the amygdala in fear conditioning measured with fear-potentiated startle in rats trained concurrently with auditory and visual conditioned stimuli. The Journal of Neuroscience 15:2312–2327.

Cenquizca LA, Swanson LW (2007) Spatial organization of direct hippocampal field CA1 axonal projections to the rest of the cerebral cortex. Brain Res Rev 56:1–26.

Chen AC, Etkin A (2013) Hippocampal network connectivity and activation differentiates post-traumatic stress disorder from generalized anxiety disorder. Neuropsychopharmacology 38:1889–1898.

Cullen PK, Gilman TL, Winiecki P, Riccio DC, Jasnow AM (2015) Activity of the anterior cingulate cortex and ventral hippocampus underlie increases in contextual fear generalization. NEUROBIOLOGY OF LEARNING AND MEMORY 124:19–27.

Do-Monte FH, Quiñones-Laracuente K, Quirk GJ (2016) A temporal shift in the circuits mediating retrieval of fear memory. Nature 519:460–463.

Dymond S, Dunsmoor JE, Vervliet B, Roche B, Hermans D (2015) Fear Generalization in Humans: Systematic Review and Implications for Anxiety Disorder Research. Behav Ther 46:561–582.

Einarsson EÖ, Nader K (2012) Involvement of the anterior cingulate cortex in formation, consolidation, and reconsolidation of recent and remote contextual fear memory. Learning & Memory 19:449–452.

Einarsson EÖ, Pors J, Nader K (2015) Systems Reconsolidation Reveals a Selective Role for the Anterior Cingulate Cortex in Generalized Contextual Fear Memory Expression. Neuropsychopharmacology 40:480–487.

Fanselow MS, Dong H-W (2010) Are the dorsal and ventral hippocampus functionally distinct structures? Neuron 65:7–19.

Frankland PW, Bontempi B, Talton LE, Kaczmarek L, Silva AJ (2004a) The involvement of the anterior cingulate cortex in remote contextual fear memory. Science 304:881–883.

Frankland PW, Cestari V, Filipkowski RK, McDonald RJ, Silva AJ (1998) The dorsal hippocampus is essential for context discrimination but not for contextual conditioning. Behav Neurosci 112:863–874.

Frankland PW, Josselyn SA, Anagnostaras SG, Kogan JH, Takahashi E, Silva AJ (2004b) Consolidation of CS and US representations in associative fear conditioning. Hippocampus 14:557–569.

Gomez JL, Bonaventura J, Lesniak W, Mathews WB, Sysa-Shah P, Rodriguez LA, Ellis RJ, Richie CT, Harvey BK, Dannals RF, Pomper MG, Bonci A, Michaelides M (2017) Chemogenetics revealed: DREADD occupancy and activation via converted clozapine. Science 357:503–507.

Goshen I, Brodsky M, Prakash R, Wallace J, Gradinaru V, Ramakrishnan C, Deisseroth K (2011) Dynamics of retrieval strategies for remote memories. Cell 147:678–689.

Greenberg T, Carlson JM, Cha J, Hajcak G, Mujica-Parodi LR (2013) Ventromedial prefrontal cortex reactivity is altered in generalized anxiety disorder during fear generalization. Depress Anxiety 30:242–250.

Gurvits TV, Shenton ME, Hokama H, Ohta H, Lasko NB, Gilbertson MW, Orr SP, Kikinis R, Jolesz FA, McCarley RW, Pitman RK (1996) Magnetic resonance imaging study of hippocampal volume in chronic, combat-related posttraumatic stress disorder. Biological Psychiatry 40:1091–1099.

Hardt O, Nader K, Nadel L (2013) Decay happens: the role of active forgetting in memory. Trends in Cognitive Sciences 17:109–118.

Hefner K, Whittle N, Juhasz J, Norcross M, Karlsson R-M, Saksida LM, Bussey TJ, Singewald N, Holmes A (2008) Impaired fear extinction learning and cortico-amygdala circuit abnormalities in a common genetic mouse strain. J Neurosci 28:8074–8085.

Huff ML, Emmons EB, Narayanan NS, LaLumiere RT (2016) Basolateral amygdala projections to ventral hippocampus modulate the consolidation of footshock, but not contextual, learning in rats. Learning & Memory 23:51–60.

Hunsaker MR, Kesner RP (2008) Dissociations across the dorsal-ventral axis of CA3 and CA1 for encoding and retrieval of contextual and auditory-cued fear. NEUROBIOLOGY OF LEARNING AND MEMORY 89:61–69.

Jasnow AM, Cullen PK, Riccio DC (2012) Remembering another aspect of forgetting. Front Psychol 3:175.

Jasnow AM, Lynch JF III, Gilman TL, Riccio DC (2016) Perspectives on fear generalization and its implications for emotional disorders. Journal of Neuroscience Research 95:821–835.

Jendryka M, Palchaudhuri M, Ursu D, van der Veen B, Liss B, Kätzel D, Nissen W, Pekcec A (2019) Pharmacokinetic and pharmacodynamic actions of clozapine-N-oxide, clozapine, and compound 21 in DREADD-based chemogenetics in mice. Sci Rep 9:4522.

Kessler RC, Petukhova M, Sampson NA, Zaslavsky AM, Wittchen H-U (2012) Twelve-month and lifetime prevalence and lifetime morbid risk of anxiety and mood disorders in the United States. international journal of methods in psychiatric research 21:169–184.

Kim JJ, Fanselow MS (1992) Modality-specific retrograde amnesia of fear. Science 256:675–677.

Kim JJ, Rison RA, Fanselow MS (1993) Effects of amygdala, hippocampus, and periaqueductal gray lesions on short- and long-term contextual fear. Behav Neurosci 107:1093–1098.

Lissek S, Powers AS, McClure EB, Phelps EA, Woldehawariat G, Grillon C, Pine DS (2005) Classical fear conditioning in the anxiety disorders: a meta-analysis. Behav Res Ther 43:1391–1424.

Lissek S, Rabin S, Heller RE, Lukenbaugh D, Geraci M, Pine DS, Grillon C (2010) Overgeneralization of conditioned fear as a pathogenic marker of panic disorder. Am J Psychiatry 167:47–55.

Lynch JF, Winiecki P, Gilman TL, Adkins JM, Jasnow AM (2017) Hippocampal GABAB(1a) Receptors Constrain Generalized Contextual Fear. Neuropsychopharmacology 42:914–924.

MacLaren DAA, Browne RW, Shaw JK, Krishnan Radhakrishnan S, Khare P, España RA, Clark SD (2016) Clozapine N-Oxide Administration Produces Behavioral Effects in Long-Evans Rats: Implications for Designing DREADD Experiments. eNeuro 3.

Mahler SV, Vazey EM, Beckley JT, Keistler CR, McGlinchey EM, Kaufling J, Wilson SP, Deisseroth K, Woodward JJ, Aston-Jones G (2014) Designer receptors show role for ventral pallidum input to ventral tegmental area in cocaine seeking. Nat Neurosci 17:577–585.

Manvich DF, Webster KA, Foster SL, Farrell MS, Ritchie JC, Porter JH, Weinshenker D (2018) The DREADD agonist clozapine N-oxide (CNO) is reverse-metabolized to clozapine and produces clozapine-like interoceptive stimulus effects in rats and mice. Sci Rep 8:3840.

Maren S, Aharonov G, Stote DL, Fanselow MS (1996) N-methyl-D-aspartate receptors in the basolateral amygdala are required for both acquisition and expression of conditional fear in rats. Behav Neurosci 110:1365–1374.

Maren S, Fanselow MS (1995) Synaptic plasticity in the basolateral amygdala induced by hippocampal formation stimulation in vivo. Journal of Neuroscience 15:7548–7564.

Maren S, Holt WG (2004) Hippocampus and Pavlovian fear conditioning in rats: muscimol infusions into the ventral, but not dorsal, hippocampus impair the acquisition of conditional freezing to an auditory conditional stimulus. Behav Neurosci 118:97–110.

McCullough KM, Morrison FG, Ressler KJ (2016) Bridging the Gap: Towards a cell-type specific understanding of neural circuits underlying fear behaviors. NEUROBIOLOGY OF LEARNING AND MEMORY 135:27–39.

Morey RA, Dunsmoor JE, Haswell CC, Brown VM, Vora A, Weiner J, Stjepanovic D, Wagner HR, VA Mid-Atlantic MIRECC Workgroup, LaBar KS (2015) Fear learning circuitry is biased toward generalization of fear associations in posttraumatic stress disorder. Transl Psychiatry 5:e700–e700.

Morozov A, Sukato D, Ito W (2011) Selective Suppression of Plasticity in Amygdala Inputs from Temporal Association Cortex by the External Capsule. Journal of Neuroscience 31:339–345.

Morris RGM (1981) Spatial localization does not require the presence of local cues. Learn Motiv 12:239–260.

Nadel L, Moscovitch M (1997) Memory consolidation, retrograde amnesia and the hippocampal complex. Curr Opin Neurobiol 7:217–227.

Schafe GE, Doyère V, LeDoux JE (2005) Tracking the fear engram: the lateral amygdala is an essential locus of fear memory storage. J Neurosci 25:10010–10014.

Schuff N, Neylan TC, Lenoci MA, Du AT, Weiss DS, Marmar CR, Weiner MW (2001) Decreased hippocampal N-acetylaspartate in the absence of atrophy in posttraumatic stress disorder. Biological Psychiatry 50:952–959.

Scofield MD, Boger HA, Smith RJ, Li H, Haydon PG, Kalivas PW (2015) Gq-DREADD Selectively Initiates Glial Glutamate Release and Inhibits Cue-induced Cocaine Seeking. Biological Psychiatry 78:441–451.

Shin LM, Orr SP, Carson MA, Rauch SL, Macklin ML, Lasko NB, Peters PM, Metzger LJ, Dougherty DD, Cannistraro PA, Alpert NM, Fischman AJ, Pitman RK (2004) Regional cerebral blood flow in the amygdala and medial prefrontal cortex during traumatic imagery in male and female Vietnam veterans with PTSD. Arch Gen Psychiatry 61:168–176.

Shin LM, Rauch SL, Pitman RK (2006) Amygdala, medial prefrontal cortex, and hippocampal function in PTSD. Ann N Y Acad Sci 1071:67–79.

Smith DR, Gallagher M, Stanton ME (2007) Genetic background differences and nonassociative effects in mouse trace fear conditioning. Learn Mem 14:597–605.

Squire LR, Alvarez P (1995a) Retrograde amnesia and memory consolidation: a neurobiological perspective. Curr Opin Neurobiol 5:169–177.

Squire LR, Alvarez P (1995b) Retrograde amnesia and memory consolidation: a neurobiological perspective. Curr Opin Neurobiol 5:169–177.

Stachniak TJ, Ghosh A, Sternson SM (2014) Chemogenetic synaptic silencing of neural circuits localizes a hypothalamus→midbrain pathway for feeding behavior. Neuron 82:797–808.

Tanaka KZ, Pevzner A, Hamidi AB, Nakazawa Y, Graham J, Wiltgen BJ (2014) Cortical Representations Are Reinstated by the Hippocampus during Memory Retrieval. Neuron 84:347–354.

Teyler TJ, Rudy JW (2007) The hippocampal indexing theory and episodic memory: Updating the index. Hippocampus 17:1158–1169.

Vazey EM, Aston-Jones G (2014) Designer receptor manipulations reveal a role of the locus coeruleus noradrenergic system in isoflurane general anesthesia. Proceedings of the National Academy of Sciences 111:3859–3864.

Whissell PD, Tohyama S, Martin LJ (2016) The Use of DREADDs to Deconstruct Behavior. Front Genet 7:70.

Wiltgen BJ, Silva AJ (2007) Memory for context becomes less specific with time. Learn Mem 14:313–317.

Wiltgen BJ, Zhou M, Cai Y, Balaji J, Karlsson MG, Parivash SN, Li W, Silva AJ (2010) The Hippocampus Plays a Selective Role in the Retrieval of Detailed Contextual Memories. Current Biology 20:1336–1344.

Winocur G, Frankland PW, Sekeres M, Fogel S, Moscovitch M (2009) Changes in context-specificity during memory reconsolidation: selective effects of hippocampal lesions. Learn Mem 16:722–729.

Winocur G, Moscovitch M, Sekeres M (2007) Memory consolidation or transformation: context manipulation and hippocampal representations of memory. Nat Neurosci 10:555–557.

Winocur G, Sekeres MJ, Binns MA, Moscovitch M (2013) Hippocampal lesions produce both nongraded and temporally graded retrograde amnesia in the same rat. Hippocampus 23:330–341.

Woodward SH, Kaloupek DG, Streeter CC, Martinez C, Schaer M, Eliez S (2006) Decreased Anterior Cingulate Volume in Combat-Related PTSD. Biological Psychiatry 59:582–587.

Xu W, Südhof TC (2013) A Neural Circuit for Memory Specificity and Generalization. Science 339:1290–1295.

Yamasue H, Kasai K, Iwanami A, Ohtani T, Yamada H, Abe O, Kuroki N, Fukuda R, Tochigi M, Furukawa S, Sadamatsu M, Sasaki T, Aoki S, Ohtomo K, Asukai N, Kato N (2003) Voxel-based analysis of MRI reveals anterior cingulate gray-matter volume reduction in posttraumatic stress disorder due to terrorism. Proceedings of the National Academy of Sciences 100:9039–9043.

Zhou H, Xiong G-J, Jing L, Song N-N, Pu D-L, Tang X, He X-B, Xu F-Q, Huang J-F, Li L-J, Richter-Levin G, Mao R-R, Zhou Q-X, Ding Y-Q, Xu L (2017) The interhemispheric CA1 circuit governs rapid generalisation but not fear memory. Nat Commun 8:2190.

Zola-Morgan SM, Squire LR (1990) The primate hippocampal formation: evidence for a time-limited role in memory storage. Science 250:288–290.

